# An egg sabotaging mechanism drives non-Mendelian transmission in mice

**DOI:** 10.1101/2024.02.22.581453

**Authors:** Frances E. Clark, Naomi L. Greenberg, Duilio M.Z.A. Silva, Emily Trimm, Morgan Skinner, R Zaak Walton, Leah F. Rosin, Michael A. Lampson, Takashi Akera

## Abstract

During meiosis, homologous chromosomes segregate so that alleles are transmitted equally to haploid gametes, following Mendel’s Law of Segregation. However, some selfish genetic elements drive in meiosis to distort the transmission ratio and increase their representation in gametes. The established paradigms for drive are fundamentally different for female vs male meiosis. In male meiosis, selfish elements typically kill gametes that do not contain them. In female meiosis, killing is predetermined, and selfish elements bias their segregation to the single surviving gamete (i.e., the egg in animal meiosis). Here we show that a selfish element on mouse chromosome 2, *R2d2*, drives using a hybrid mechanism in female meiosis, incorporating elements of both male and female drivers. If *R2d2* is destined for the polar body, it manipulates segregation to sabotage the egg by causing aneuploidy that is subsequently lethal in the embryo, so that surviving progeny preferentially contain *R2d2*. In heterozygous females, *R2d2* orients randomly on the metaphase spindle but lags during anaphase and preferentially remains in the egg, regardless of its initial orientation. Thus, the egg genotype is either euploid with *R2d2* or aneuploid with both homologs of chromosome 2, with only the former generating viable embryos. Consistent with this model, *R2d2* heterozygous females produce eggs with increased aneuploidy for chromosome 2, increased embryonic lethality, and increased transmission of *R2d2*. In contrast to a male meiotic driver, which kills its sister gametes produced as daughter cells in the same meiosis, *R2d2* eliminates “cousins” produced from meioses in which it should have been excluded from the egg.

## Introduction

Mendel’s Law of Segregation states that each pair of allele segregates randomly during meiosis, transmitting to gametes at an equal (50%) chance. However, some genetic elements violate this law to preferentially transmit to the next generation with associated fitness costs to the host (e.g., reduced fertility).^1–7^ Such selfish elements that cheat during meiosis are called meiotic drivers, and distinct mechanisms to achieve their biased transmission have been identified between male and female meiosis.^8^ Female meiotic drive is thought to be accomplished through preferential segregation to the egg, exploiting the inherent asymmetry in female meiosis: only the chromosomes that segregate to the egg will transmit.^3,9–11^ In contrast, male meiosis undergoes symmetric divisions, and cheating mostly happens post-meiosis.^5,7,8^ Male (and spore killer) meiotic drivers typically act through the sabotage or death of competitor sperm/spores that do not carry them.^5,7,12^ Although disabling non-carrier sperm often reduces fertility, sperm killer systems allow more eggs to be fertilized by sperm that carry meiotic drivers, leading to biased transmission to the offspring.^13^

Female meiotic drivers manipulate chromosome segregation to remain in the egg. Centromeres have an ideal opportunity to cheat in female meiosis because they interact with the spindle to direct chromosome orientation and segregation.^14–19^ Indeed, selfish expanded centromeres preferentially remain in the egg in both animals and plants.^20–22^ In animals, selfish mouse centromeres have been the primary system to study cell biological mechanisms underlying female meiotic drive.^21,23–25^ Other loci usually do not control chromosome-spindle interactions, and therefore, it is unknown how non-centromeric meiotic drivers manipulate their segregation patterns, except for maize knob that exploit an asymmetry specific to plant female meiosis.^3,26–28^ We chose the selfish *R2d2* (*Responder to drive 2*) locus as a model system to tackle this question. *R2d2* is a repetitive DNA of a 127 kb-long monomer found on mouse chromosome 2 and experiences transmission rates above 95% from heterozygous female mice.^29–31^ Transmission from heterozygous male mice is Mendelian (i.e., 50%).^29^ The female meiotic drive is associated with a fertility cost of ~30% reduction in litter size due to embryonic lethality.^29^ The molecular mechanisms underlying biased transmission and embryonic lethality are completely unknown.

## Results

### Cell biological approaches to investigate *R2d2* meiotic drive

*R2d2* is located in the middle of a telocentric chromosome arm (i.e., Ch.2: 83,790,939 – 84,701,151 in the GRCm38/mm10 mouse reference assembly where chromosome 2 represents 182 Mb).^29^ This locus is a duplication of *R2d1*, present ~6 Mb away on the same chromosome (Figure S1A).^30^ After the initial duplication event that occurred between 2 - 3.5 million years ago, the *R2d2* locus expanded, resulting in a repetitive tandem arrayed element. Consequently, some mouse strains in the *Mus* genus completely lack *R2d2*, some possess a single copy of the *R2d2* monomer, and others have several copies of *R2d2* tandemly repeated (e.g., the WSB/EiJ strain is homozygous for 33 copies of the *R2d2* monomer).^29^

As an experimental system to study *R2d2* meiotic drive, we used *Mus musculus domesticus* intra-subspecific hybrid female mice heterozygous for the *R2d2* locus, which were previously shown to exhibit biased transmission (Figure 1A, +/–).^29^ We developed two complementary methods to visualize the *R2d2* locus in mouse oocytes: CRISPR/Cas9 based labeling using dCas9-EGFP plus gRNA targeting the *R2d2* DNA sequence and Oligopaint FISH (Figures S1 and S2, see Methods).^32,33^ Oligopaint FISH shows more robust labeling than dCas9-EGFP in fixed oocytes, while dCas9-EGFP is compatible with both fixed and live cell imaging.

**Figure 1.**
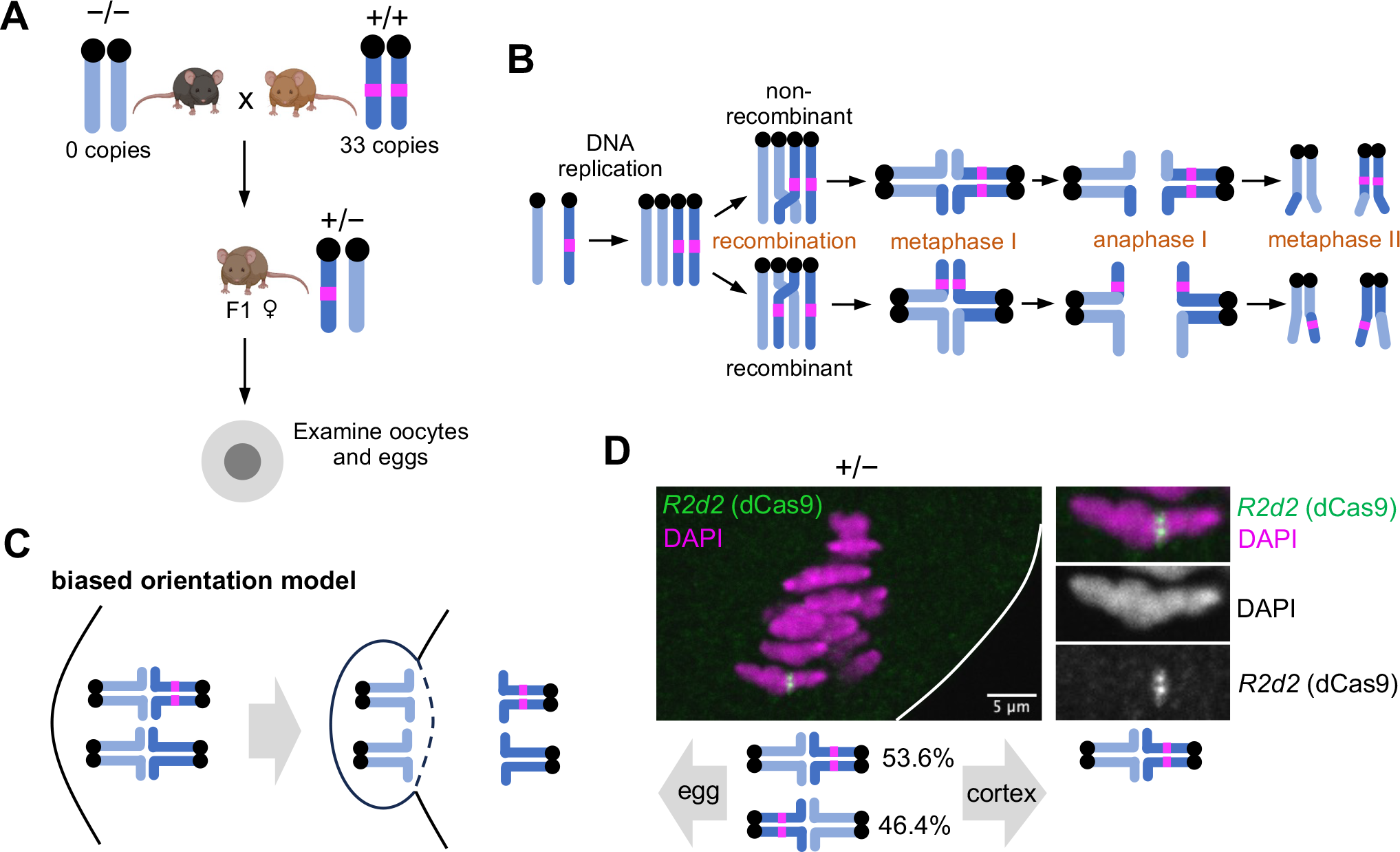
*R2d2* randomly orients on the metaphase I spindle in female meiosis. (**A**) Female F1 hybrids between two *M. m. domesticus* strains with different *R2d2* copy numbers (*R2d2* heterozygous mice, +/–) as experimental systems to study meiotic drive of *R2d2*. Oocytes and eggs collected from +/– mice (C57BL/6J x WSB/EiJ and BXD19/TyJ x WSB/EiJ hybrids) and –/– mice (the CF1 strain) were used throughout this study unless specified in the figure legend. (**B**) Schematic showing recombinant and non-recombinant chromosomes, depending on whether there is crossover between *R2d2* and the centromere. Non-recombinants can cheat in meiosis I whereas recombinants can cheat in meiosis II. (**C**) Schematic showing the biased orientation model for *R2d2* meiotic drive. (**D**) *R2d2* heterozygous (+/–) oocytes expressing dCas9-EGFP and gRNA targeting the *R2d2* sequence were fixed shortly before anaphase I and stained for EGFP. The fraction of oocytes with the *R2d2* locus oriented toward either the egg or cortical pole was measured; n = 39 cells. White line, oocyte cortex. Note that we focused only on non-recombinant chromosomes, which can cheat in meiosis I.

Female meiotic drive can occur in either of the two meiotic divisions, depending on when the driving locus segregates from its homolog.^26^ Homologous centromeres segregate in meiosis I, and therefore selfish centromeres cheat exclusively in meiosis I.^34^ In contrast, a non-centromeric locus like *R2d2* can cheat either in meiosis I or II depending on the crossover position (Figure 1B).^27,35^ If the crossover is not located between *R2d2* and the centromere (non-recombinant), one pair of sister chromatids has *R2d2* and the other pair does not (Figure 1B, top). In this case, the *R2d2* locus segregates from its homolog in meiosis I, and the cheating happens in meiosis I as with centromeres. Alternatively, crossover between the centromere and *R2d2* will create recombinant chromatids, where one sister chromatid of each pair has *R2d2* and the other sister does not (Figure 1B, bottom). In this configuration, both products of meiosis I receive one chromatid with *R2d2*, so there is no opportunity for cheating in meiosis I. *R2d2* segregates from its homolog in meiosis II, providing the opportunity to bias its segregation to the egg. Given *R2d2*’s central location within the chromosome arm, it could in principle cheat in either meiosis I or meiosis II, depending on the crossover position.

### *R2d2* randomly orients on the metaphase I spindle in female meiosis

Selfish centromeres preferentially orient towards the interior side of the metaphase I spindle to preferentially remain the egg.^20,21^ Depending on the crossover position, *R2d2* could cheat by this mechanism in either metaphase I (non-recombinant) or metaphase II (recombinant). We tested for biased orientation in metaphase I (Figure 1C), when the spindle is oriented perpendicular to the cell cortex, so that we can determine which spindle pole will be extruded to the polar body and which will be retained in the egg.^36^ In contrast, the spindle is parallel to the cortex at metaphase II, so we cannot predict which pole will be extruded to the polar body.^37,38^ Using dCas9-EGFP to visualize *R2d2*, we did not find a significant orientation bias of non-recombinant chromosomes in metaphase I (Figure 1D). Based on this finding, it is unlikely that *R2d2* biases its transmission by biased orientation on the spindle, indicating that *R2d2* and centromeres use distinct cheating strategies.

### Anaphase lagging model for meiotic drive

Another possible cheating mechanism for *R2d2* that is compatible with both meiosis I and II is to induce its own chromosome to lag in anaphase, without affecting anaphase movement of the paired chromosome 2 lacking *R2d2*. Because of the highly asymmetric cell division in female meiosis, lagging chromosomes can be carried with the larger amount of cytoplasm into the egg (Figure 2A). If *R2d2* initially orients towards the interior side of the meiosis I spindle, it lags and still ends up in the egg, with the homologous chromosome 2 in the polar body (Figure 2Ai). If *R2d2* initially orients towards the cortex, it lags and remains in the egg along with the homologous chromosome 2, causing aneuploidy of chromosome 2 and subsequent embryonic lethality (Figure 2Aii).^39,40^ Similar mechanisms can be applied to meiosis II where recombinant sister chromatids with and without *R2d2* compete for segregation to the egg. This embryo killing strategy leads to biased transmission of *R2d2* because all surviving embryos are euploid with *R2d2*.

**Figure 2.**
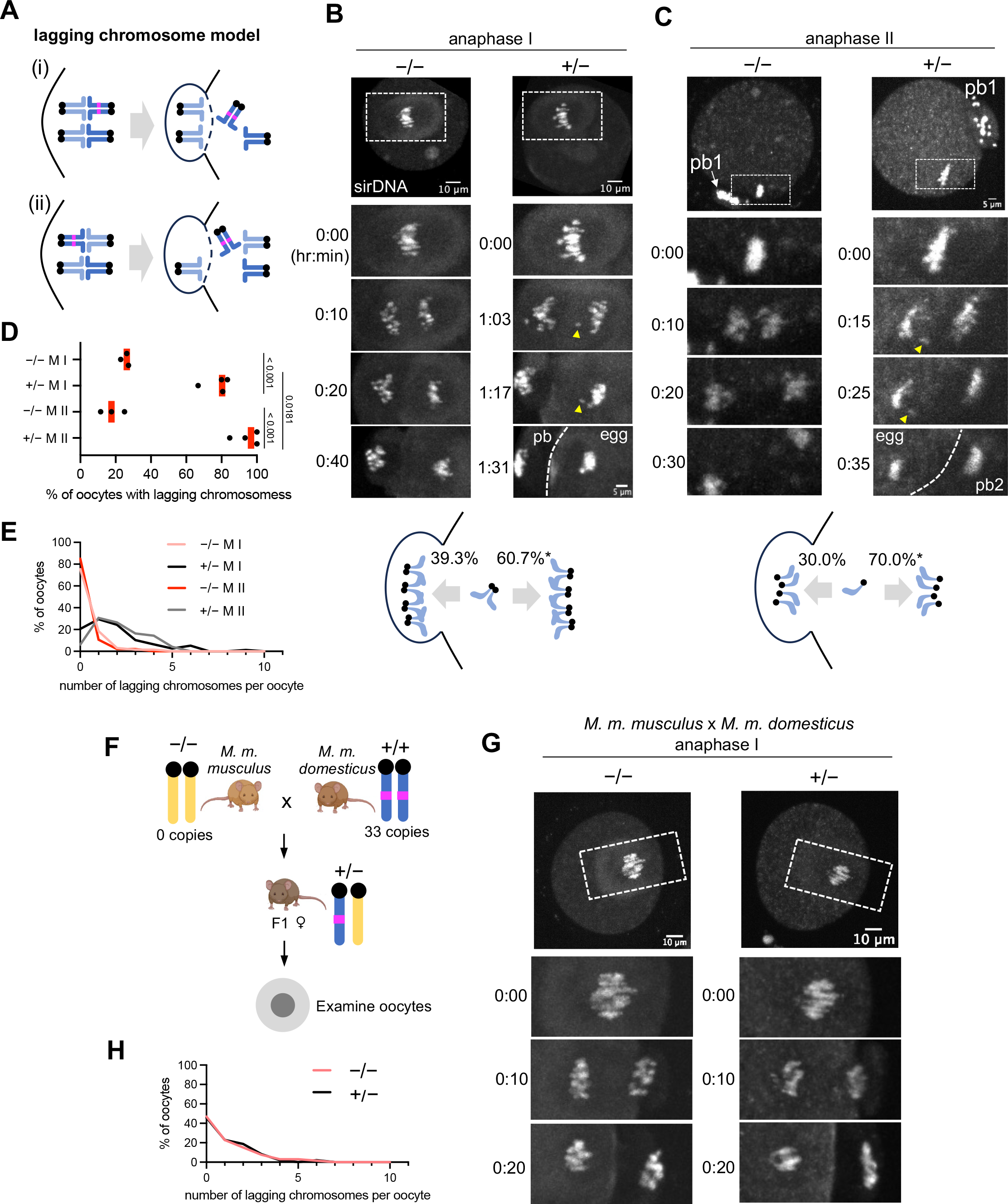
*R2d2* meiotic drive is associated with increased anaphase lagging. (**A**) Schematic showing the anaphase lagging model for *R2d2* meiotic drive. (**B, C**) Control (–/–) and *R2d2* heterozygous (+/–) meiosis I oocytes (B) and meiosis II eggs (C) were stained with sirDNA to visualize chromosomes and imaged live at anaphase (n = 65 and 80 meiosis I oocytes for control and *R2d2* heterozygote, respectively and n = 47 and 49 meiosis II eggs for control and *R2d2* heterozygote, respectively). Note that the lagging chromosomes (yellow arrowheads) eventually remained in the egg. The segregation pattern of lagging chromosomes was quantified; chi-square test for goodness of fit was used to examine the deviation from the expected 50:50 ratio (B, n = 145 lagging chromosomes from 76 meiosis I oocytes; \**P* = 0.01004 and C, n = 40 lagging chromosomes from 17 meiosis II eggs; \**P* = 0.01141). (**D, E**) Based on the live imaging data in B and C, the fraction of oocytes with at least one lagging chromosome (D, each dot represents an independent experiment; red line, mean; unpaired two-tailed t-test was performed) and the distribution of the number of lagging chromosomes per oocyte were quantified. (**F**) Female F1 hybrids between a *M. m. domesticus* strain with expanded *R2d2* (WSB/EiJ) and a *M. m. musculus* strain without it (PWD/PhJ) show Mendelian segregation of *R2d2*. (**G, H**) Control (–/–, PWD/PhJ x C57BL/6J) and *R2d2* heterozygous (+/–, PWD/PhJ x WSB/EiJ) oocytes in the *M. m. domesticus* x *M. m. musculus* hybrid genetic background were stained with sirDNA to visualize chromosomes and imaged live at anaphase I (G), and the distribution of the number of lagging chromosomes per oocyte was quantified (H, n = 66 and 48 oocytes for control and *R2d2* heterozygote, respectively).

The anaphase lagging model makes several predictions that can be experimentally tested: (1) lagging chromosomes should be present at high frequency in crosses where cheating occurs, (2) lagging chromosomes preferentially end up in the egg rather than the polar body, (3) chromosome 2 with *R2d2* lags more frequently than other chromosomes, and (4) aneuploidy for chromosome 2 is more frequent than for other chromosomes in crosses where cheating occurs. Although these predictions apply to both meiosis I and meiosis II, depending on the crossover position, we focused primarily on meiosis I because of technical difficulty in efficiently labeling *R2d2* in anaphase II.

We tested the first and second predictions by live imaging of *R2d2* heterozygous oocytes and control oocytes without *R2d2*. Heterozygous oocytes had significantly more lagging chromosome events in anaphase I and II compared to controls (Figures 2B-2E). Furthermore, these lagging chromosomes preferentially remained in the egg (Figures 2B and 2C). To test whether anaphase lagging is functionally related to preferential transmission, we examined a hybrid that is heterozygous for *R2d2* but does not exhibit meiotic drive. Previous studies have revealed that the strength of *R2d2* drive differs depending on the strains involved (reporting transmission ratios ranging from 50% to 95%), which suggests that other unlinked loci modify the strength of drive.^29^ For example, no drive was observed when the same *M. m. domesticus* strain with expanded *R2d2* used in our previous experiments was crossed with a *M. m. musculus* strain (Figure 2F). Oocytes from this non-driving hybrid did not show an increased anaphase lagging rate compared to the control *M. m. musculus* x *M. m. domesticus* hybrid without *R2d2* (Figures 2G and 2H). Together these observations are consistent with the first two predictions of the anaphase lagging model for biased transmission of *R2d2*.

To test the third prediction, we visualized *R2d2* in anaphase I using dCas9-EGFP. We found that 63.9% of heterozygous oocytes experienced chromosome 2 with *R2d2* lagging in anaphase I while other chromosomes segregated to the spindle poles (n = 36 oocytes) (Figure 3A). In some oocytes, we were able to distinguish between recombinant and non-recombinant chromosomes and found anaphase I lagging in both cases. Only non-recombinant chromosomes can cheat in meiosis I, but most of the recombinant chromosomes appear to eventually segregate equally in meiosis I, allowing them to cheat in meiosis II (see below).

**Figure 3.**
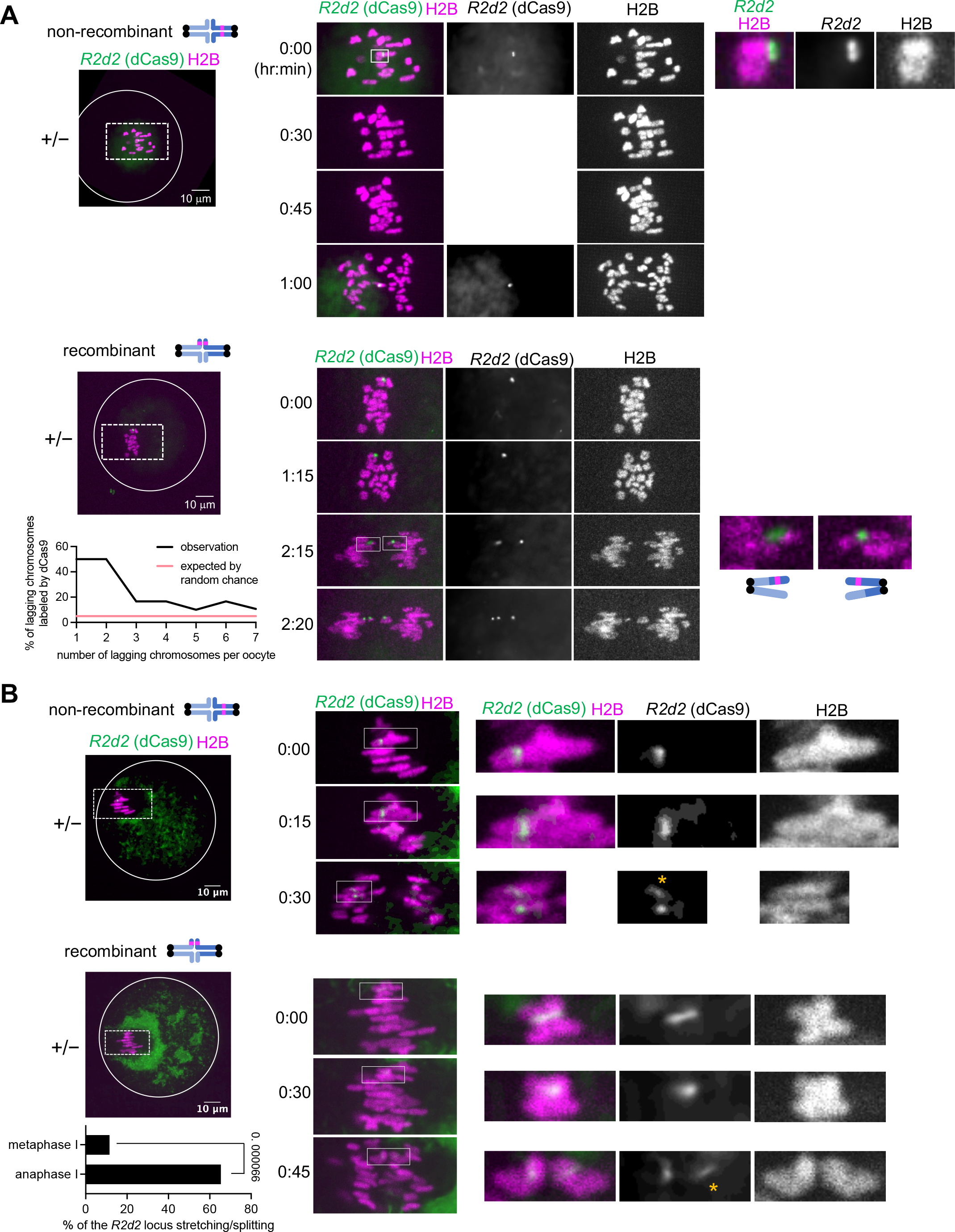
*R2d2* induces its own chromosome to lag in anaphase. (**A**) *R2d2* heterozygous (+/–) oocytes expressing H2B-mCherry, dCas9-EGFP, and gRNA targeting the *R2d2* DNA sequence were imaged live at anaphase I (white line, oocyte cortex). The histogram shows the distribution of the fraction of lagging chromosomes that have dCas9 signals (n = 35 cells). For the non-recombinant chromosome example (top), we imaged the dCas9-EGFP signals only at the first timepoint and at the anaphase onset to minimize photobleaching and phototoxicity. In the recombinant chromosome example (bottom), three dCas9 signals were observed at the 2 hr 20 min timepoint due to the splitting of the *R2d2* locus on one of the homologs (also see B). (**B**) *R2d2* heterozygous (+/–) oocytes expressing H2B-mCherry, dCas9-EGFP, and gRNA targeting the *R2d2* DNA sequence were imaged live at the metaphase I to anaphase I transition. The fraction of oocytes with the dCas9 signal splitting or stretching was quantified in metaphase I and anaphase I (n = 26 cells); Chi-Square Test of Independence was used to calculate the *P* values in the graph; white line, oocyte cortex; orange asterisks, stretching *R2d2* locus. We used the Denoise.ai software (Nikon) to reduce noise and follow the *R2d2* locus better in anaphase I.

Given that the mouse karyotype is 2n = 40, if chromosome 2 with *R2d2* lags at the same rate as other chromosomes, we expect to see one in 20 lagging chromosomes (i.e., 5%) with dCas9 labeling regardless of how many lagging chromosomes are present in each oocyte (Figure 3A graph, pink line). Instead, we found that chromosome 2 with *R2d2* was lagging more often than expected by random chance (Figure 3A graph, black line). Moreover, the fraction of lagging chromosomes with *R2d2* was especially high when there were only one or two lagging chromosomes in anaphase I, as expected if most lagging events are due to *R2d2*. These findings are consistent with the third model prediction and suggest that *R2d2* can induce its own chromosome to lag in anaphase as a mechanism to bias its transmission to surviving progeny.

To induce lagging, *R2d2* likely interacts with some intracellular structure to slow its poleward movement during anaphase. If the centromere of chromosome 2 is pulled towards the spindle pole but the *R2d2* locus is resisting poleward movement, the *R2d2* locus should experience tension due to the tug-of-war. Consistent with this idea, we often observed splitting and stretching of the *R2d2* locus in anaphase I (Figures 3A, recombinant, and 3B), implying that *R2d2* is interacting with some structure to induce lagging. As a possible alternative explanation for anaphase I lagging, bivalents containing *R2d2* might fail to resolve into univalents. Live imaging of centromeres in *R2d2* heterozygous oocytes (Figure S3) revealed that 96.8% of the anaphase lagging chromosomes separated properly into univalent chromosomes, arguing against this possibility.

To test the fourth prediction regarding the higher aneuploidy for chromosome 2 with *R2d2*, we measured aneuploidy in metaphase II eggs using Oligopaint FISH (Figures 4A and 4B). Although the overall aneuploidy rates were low, chromosome 2 with *R2d2* showed higher aneuploidy than chromosome 1, which has a similar size (Figure 4A, *in vitro* +/– and *in vivo* +/–). The low chromosome 2 aneuploidy rate suggests that meiotic drive by anaphase lagging is weaker in meiosis I compared to meiosis II. Indeed, our results show that frequencies of lagging chromosome events and lagging chromosomes remaining in the egg are both higher in meiosis II compared to meiosis I (Figures 2B-2D). A high transmission rate would therefore depend on recombinant chromosomes cheating in meiosis II. We find by Oligopaint FISH analyses that 77.5% of metaphase II eggs contained recombinant chromatids (Figure S4), consistent with *R2d2* cheating more often in meiosis II compared to meiosis I. Even though recombinant chromosomes lag in anaphase I (Figure 3A, recombinant), they eventually segregate equally in most cases, providing the opportunity for them to cheat in meiosis II.

**Figure 4.**
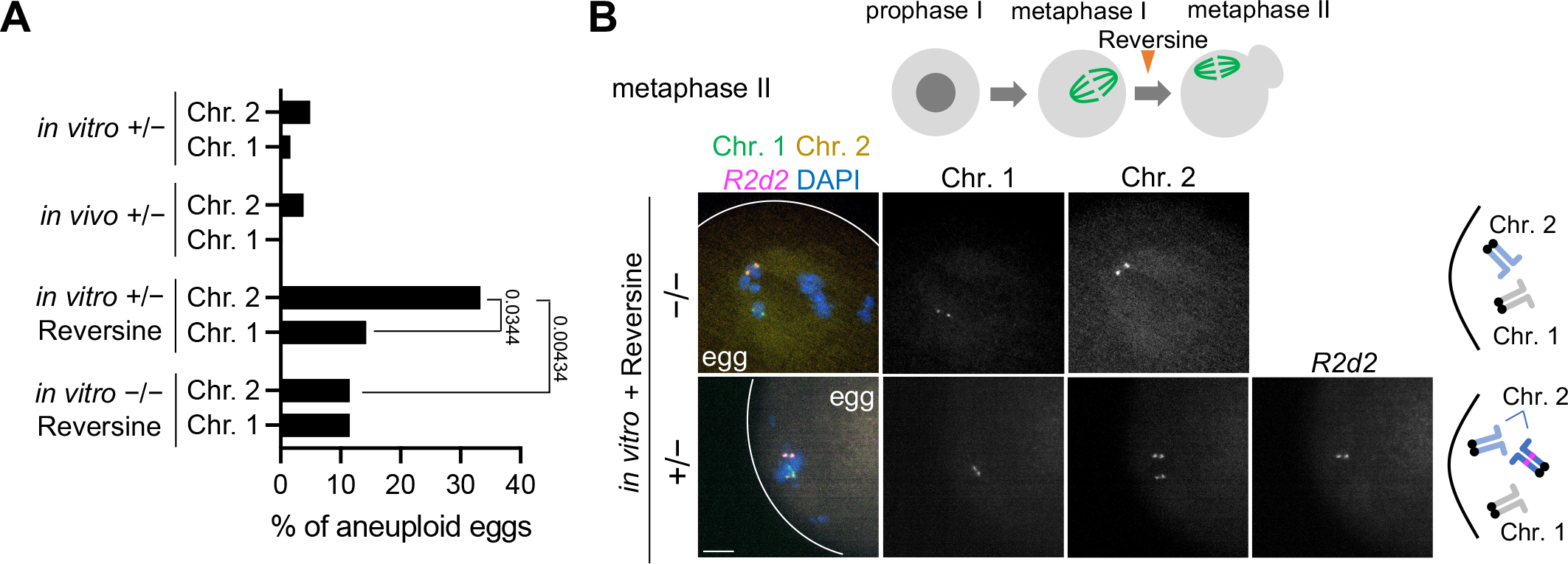
Chromosome 2 with *R2d2* has a higher aneuploid rate. (**A**) Quantification of the fraction of the oocytes that are aneuploid for chromosome 1 or 2 at the metaphase II stage (n = 52, 52, 84, 35, 162, 61, 78, and 36 oocytes for *in vitro* –/– Reversine Chr. 2, *in vitro* –/– Reversine Chr. 1, *in vitro* +/– Reversine Chr. 2, *in vitro* +/– Reversine Chr. 1, *in vitro* +/– Chr. 2, *in vitro* +/– Chr. 1, *in vivo* +/– Chr. 2, and *in vivo* +/– Chr. 1, Chi-Square Test of Independence was used to calculate the *P* values in the graph). (**B**) Top schematic shows the Reversine treatment of mouse oocytes at metaphase I. Control (–/–) and *R2d2* heterozygous (+/–) oocytes were matured *in vitro* in the presence of Reversine and fixed at metaphase II to visualize chromosome 1 and 2 and the *R2d2* locus with Oligopaint FISH. White lines, egg cell cortex.

Because of the large number of animals required to compare low aneuploidy rates, we sensitized the system by inhibiting a spindle checkpoint kinase, MPS1, with the inhibitor Reversine to increase the basal aneuploidy levels. We collected prophase I oocytes from *R2d2* heterozygous mice, treated with Reversine at metaphase I, and matured them *in vitro* until metaphase II (Figure 4B). Reversine-treated eggs confirmed our finding that chromosome 2 with *R2d2* has a higher aneuploidy rate than chromosome 1 (Figures 4A and 4B). To test whether the presence of *R2d2* is involved in the higher aneuploidy rate of chromosome 2, oocytes from wildtype (–/–) mice were matured to metaphase II *in vitro* in the presence of Reversine. The aneuploid rate of chromosome 2 without *R2d2* was similar to that of chromosome 1 and significantly less compared to chromosome 2 with *R2d2* (Figures 4A and 4B). Collectively, these results are consistent with our anaphase lagging model where *R2d2* can increase chromosome 2 aneuploidy via lagging to bias its transmission (Figure 2A).

## Discussion

### *R2d2* sabotages the egg with the “wrong” genotype to bias its transmission

The anaphase lagging model incorporates elements of both female and male meiotic drive systems. Female meiotic drivers typically manipulate segregation to preferentially remain in the egg, without creating aneuploidy.^3,41,42^ *R2d2* also manipulates segregation, but by lagging in anaphase to create aneuploidy instead of segregating to the polar body. Male (and spore killer) meiotic drivers typically act through the sabotage or death of competitor gametes.^43–46^ Similarly, the aneuploidy induced by *R2d2* lagging is expected to lead to embryonic lethality. This mechanism does not increase the absolute number of euploid eggs possessing *R2d2* but instead enriches *R2d2* among surviving offspring. This model is consistent with two previous findings.^29,47^ First, stronger *R2d2* meiotic drive is associated with higher embryonic lethality rates as a fitness cost. Second, *R2d2* biases its transmission rate mainly by decreasing the absolute number of wild-type progeny without *R2d2* rather than increasing the number of progeny with *R2d2*. Therefore, the anaphase lagging model can explain the mechanisms underlying both the biased transmission and the fitness cost associated with *R2d2* meiotic drive. This work provides cell biological insights into how non-centromeric loci can bias their transmission through female meiosis and raises two important questions. First, how does *R2d2* induce anaphase lagging? Second, how do lagging chromosomes remain in the egg?

### How does *R2d2* induce anaphase lagging and remain in the egg?

In some plants, accessory B chromosomes induce anaphase lagging by preventing chromosome separation to bias their segregation during pollen mitosis.^48–50^ In contrast, we suspect that *R2d2* must interact with some structures like the spindle to maintain its position during anaphase to induce lagging. This idea is supported by the observations that chromosomes separate normally in anaphase (Figure S3) and that the *R2d2* locus is under tension in anaphase (Figure 3B). One possibility is that *R2d2* interacts with the spindle using microtubule-binding proteins, as seen with maize knob domains, which recruit kinesin-14 motor proteins.^27,28^ Alternatively, *R2d2* may interacts with a different cellular structure such as the actin network or endomembranes surrounding the oocyte spindle.^51–53^

We found that anaphase lagging chromosomes preferentially segregated to the egg in both meiosis I and II (Figures 2B and 2C). This preferential retention may be specific to the mechanism of *R2d2* lagging because of the structure with which it interacts.^52^ Alternatively, any lagging chromosomes in mouse oocytes may be preferentially retained because of the asymmetric cell division that keeps most of the cytoplasm in the egg. In this case, any mechanism that increases lagging could be sufficient for biased transmission and could potentially be exploited by other meiotic drivers in mice. Interestingly, univalent chromosomes lag and preferentially segregate to the polar body in *C. elegans*, implying species divergence in how oocytes handle anaphase lagging chromosomes.^54^

### Recombination pattern and *R2d2* meiotic drive

Roughly three quarters of metaphase II eggs from *R2d2* heterozygous mice harbored a crossover in between *R2d2* and the centromere (Figure S4), indicating that *R2d2* has the opportunity to cheat more often in meiosis II. Metaphase II eggs from the non-driving hybrid also exhibited a high rate of such recombinant chromosomes (Figure S5). These findings imply that the *R2d2* locus has an intrinsic property that induces a high recombinant rate independently of meiotic drive. If anaphase lagging is a more effective drive strategy in meiosis II, as suggested by our analyses of lagging chromosomes (Figures 2B-2D), then increased recombination may be part of a mechanism acquired by *R2d2* to further increase its ability to drive. A local decrease of recombination frequency surrounding the *R2d2* locus has been reported.^30^ However, the impact of *R2d2* repeats on the overall recombination landscape of chromosome 2 was not expected and would be interesting to pursue in both male and female meiosis in the future.

In conclusion, this work provides the first cell biological insights into how a non-centromeric selfish element can bias its transmission through female meiosis in animals. Asymmetry in cell fate is a fundamental difference between female and male meiosis, which may have originated as a strategy for selfish meiotic drivers to increase their transmission by gamete killing.^55^ Such strategies are well documented in male meiosis, and our findings indicate that they also persist in female meiosis except that *R2d2* sabotages gametes that are products of different meiotic divisions rather than sisters from the same division. Anaphase lagging chromosomes can be induced by multiple different mechanisms, including merotelic attachments at centromeres and DNA bridges on the chromosome arm. Future work will elucidate if anaphase lagging is a strategy used by other female meiotic drive systems.

### Limitations of the study

We provide evidence supporting the anaphase lagging - egg sabotaging model primarily using meiosis I oocytes, but we are unable to conduct several analyses with meiosis II eggs due to technical difficulty labeling the *R2d2* locus in anaphase II. Our anaphase lagging analyses (Figures 2C and 2D) imply that *R2d2* lags also in meiosis II, inducing aneuploidy. Alternatively, *R2d2* may employ a different cheating strategy in meiosis II to achieve the extremely strong transmission bias.

## Acknowledgements

We thank Dr. Elissa P. Lei for the comments on the manuscript, the Cell and Developmental Biology Microscopy Core Director, Dr. Andrea Stout, at the University of Pennsylvania for the assistance with the super-resolution microscopy, Fernando Pardo-Manuel de Villena, the Akera lab and the Lampson lab members for discussion, and our animal facilities for their particular care of our challenging mouse strains. This work is supported by the Intramural Programs of National Heart, Lung, and Blood Institute (1ZIAHL006249-01) (T.A.) and the *Eunice Kennedy Shriver* National Institute of Child Health and Human Development (NICHD; 1K99HD104851 to L.F.R) at the National Institutes of Health (NIH), and the NIH grant R35GM122475 (M.A.L.).

## Author Contributions

Conceptualization, T.A. and M.A.L.; Methodology, T.A., F.E.C., N.L.G., and L.F.R.; Investigation, T.A., E.T., M.S., R.Z.W., F.E.C., D.M.Z.A.S., and N.L.G.; Writing – Original Draft, F.E.C.; Writing – Review & Editing, F.E.C., N.L.G., T.A., and M.A.L.; Funding Acquisition, T.A., and M.A.L.; Resources, T.A., and M.A.L.; Supervision, T.A. and M.A.L.

## Methods

### Mouse strains

Mouse strains were purchased from Envigo (NSA, stock# 033 corresponds to CF1, *Mus musculus domesticus*), and from Jackson Laboratory (C57BL/6J, stock# 000664, *Mus musculus domesticus*, BXD19/TyJ, stock# 000010, *Mus musculus domesticus*, PWD/PhJ, Stock# 004660, *Mus musculus musculus*, WSB/EiJ, stock# 001145, *Mus musculus domesticus*). Strains CF1, C57BL/6J, BXD19/TyJ, and PWD/PhJ do not have high-copy *R2d2* repeats. WSB/EiJ has 33 copies of the *R2d2* monomer.

Reciprocal crosses were used when crossing WSB/EiJ to other strains; however, the majority of F1 females used resulted from crosses where the sire was WSB/EiJ, as it is technically challenging to cross female WSB/EiJ with other strains. F1 females from WSB/EiJ crossed to C57BL/6J or BXD19/TyJ show over 90% transmission ratio distortion of the high-copy *R2d2* repeats (Didion et al., 2015, unpublished observation by Fernando Pardo-Manuel de Villena). Mice were housed in an animal facility at room temperature, 30-70% humidity, and with a ventilated rack system. Mice were exposed to a 12 hr light/dark cycle year-round. Mice were euthanized with CO_2_ followed by cervical dislocation prior to dissection of the ovaries or ampulla. All animal experiments were approved by the Animal Care and Use Committee (National Institutes of Health Animal Study Proposal#: H-0327) and were consistent with the National Institutes of Health guidelines.

### Mouse oocyte collection and culture

All manipulation of oocytes or eggs was performed with a mouth-operated plastic pipette with either a 75, 100, or 125 µm diameter pipette tip (Cooper Surgical, Inc., Cat# MXL3-75, MXL3-100, or MXL3-125, respectively). For *in vitro*-oocyte culture, germinal vesicle-intact oocytes were collected from 6 week to 6 month-old female mice in M2 media (Sigma-Aldrich, Cat# M7167). Oocytes were then kept in a 60 x15 mm polystyrene petri dish (Falcon, cat# 351007) in either M2 media, covered with paraffin oil (Nacalai, cat# NC1506764) and incubated at 37°C in a humidified atmosphere of 0% CO_2_ in air, or transferred to M16 media (Millipore, cat# M7292), covered with paraffin oil, and incubated at 37°C in a humidified atmosphere of 5% CO_2_ in air. Oocytes were matured for variable times according to each assay. For analysis of metaphase I oocytes, cells were matured for 7 hr. For metaphase II eggs, cells were matured for 16 hr. For metaphase II eggs after the early onset of anaphase I by Reversine, cells were matured for 5 hr, transferred to M2 or M16 media with 0.5 µM Reversine (Sigma-Aldrich, Cat# R39-4-1MG), and matured for an additional 11 hr. To collect *in vivo*-ovulated oocytes, female mice were injected with 0.1 ml CARD HyperOva (Cosmo Bio USA, Cat# KYD-010-EX-X5), then 5 IU of hCG (Millipore Sigma, Cat# 9002-61-3 C1063) 48 hr later, and euthanized 17 hr after the injection to dissect out ampulla. *In vivo-*ovulated metaphase II eggs were collected from the ampulla in M2 media, then transferred to M16 media, covered with paraffin oil in a 60 x15 mm polystyrene petri dish, and incubated at 37°C in a humidified atmosphere of 5% CO_2_ in air for 1 hr.

### Oocyte microinjection

GV-intact oocytes were microinjected with ~5 pl of cRNAs or proteins in M2 containing 5 µM milrinone, using a micromanipulator TransferMan 4r and FemtoJet 4i (Eppendorf). Following the microinjection, oocytes were maintained at prophase I in M16 supplemented with 5 µM milrinone overnight to allow protein expression. cRNAs used for microinjections were *MajSat* (TALE construct that recognizes major satellite repeats fused to mClover and 3 tandem Halo tag at the C terminus) at 1500 ng/ul and *H2B-mCherry* (human histone H2B with mCherry at the C terminus) at 100 ng/µl. cRNAs were synthesized using the T7 mMessage mMachine Kit (Ambion, cat# AM1340) and purified using the MEGAclear Kit (ThermoFisher, cat# AM1908). The dCas9-EGFP-gRNA complex that targets the *R2d2* sequence was assembled *in vitro* by mixing 5 uM of dCas9-EGFP protein (Novateinbio, Cat# PR-137213G) with 5 uM of the gRNA pool for *R2d2* (see below) in the reaction buffer (2 mM HEPES, 10 mM NaCl, 5 mM MgCl2, 10 μM EDTA, pH 6.5) and incubating at room temperature for 10 min. The dCas9 complex was subsequently mixed with the H2B-mCherry cRNA and used for microinjection. The gRNA pool is a mixture of 56 gRNA that have complementary sequences to the published *R2d2* sequence.^30^ gRNA were synthesized using the GeneArt Precision gRNA Synthesis kit (Thermo Fisher, Cat# A29377). We were able to visualize the *R2d2* locus with 56 gRNA but not with 20 or 40 gRNA (data not shown). The list of primer sets that were used to produce each gRNA are provided in **Table S1**.

### Oligopaint design methods

Oligopaints were designed using a modified version of the Oligominer pipeline as previously described and the mm9 genome assembly.^33,56,57^ For the *R2d2* locus probes, the *R2d2* sequence was obtained from the Morgan et al paper.^30^ Bowtie25 was used to find oligos that uniquely mapped to the *R2d2* locus using the --very-sensitive-local alignment parameters.^58^ The final probe density was designed to be about 0.5 probes per kb, and oligos contained 42-80 bp of homology. For Oligopaints labeling the 7 Mb domain on chromosome 2, probes were at a density of 2 probes per kb with 80 bp homology. For Oligopaints labeling the middle of chromosome 1, the mm39 genome assembly was used, and probes were at a density of ~3 probes per kb with 70 bp homology. A complete list of oligo loci and sequences can be found in **Table S2**.

### Immunostaining and Oligopaint FISH of oocytes and eggs

Oocytes and eggs matured to the appropriate stage (and with the zona pellucida removed by Acidic Tyrode’s Solution (EmbryoMax, Cat# MR-004-D) for the FISH assay) were fixed in freshly prepared 2% paraformaldehyde (Thermo Scientific, Cat# 28908) in 1x PBS (Quality Biological, Cat# 119-069-101) for 20 min at room temperature (RT). Fixed cells were then washed in the blocking solution (0.3% BSA (Fisher Bioreagents, Cat# BP1600-100) and 0.01% Tween (Thermo Scientific, Cat# 9005-64-5) in 1x PBS). Cells were then permeabilized in 1x PBS with 0.1% Triton X-100 (Millipore, Cat# 9002-93-1) for 15 min at RT, then placed in the blocking solution.

For immunostaining assays, the cells were then incubated for 1 hr at RT or overnight at 4°C in the blocking solution with the goat anti-GFP antibody conjugated with Dylight488 (1:100, Rockland, cat# 600-141-215). After washed three times (10 min each) in the blocking solution, cells were then mounted on microscope slides with Antifade Mounting Medium with DAPI (Vector Laboratories, Cat# H-1200) and then sealed with L.A. Colors top coat rapid dry clear polish (Electron Microscopy Sciences, Cat# 72180).

For FISH assays, the cells were mounted on microscope slides (Fisher, cat# 12-544-3) by transferring them in a small drop of blocking solution to the surface of a 0.01% poly-L-lysine (Sigma Aldrich, Cat# 25988-63-0) coated slide. Following a 2 min incubation, the cells were washed (on the slide) with 1x PBS four times for 10 min each. Cells were then fixed a second time with 4% paraformaldehyde, 0.1% Triton X-100 in 1x PBS for 20 min at RT. The microscope slides were then washed (in Coplin jars) in 1x PBS three times for 5 min at RT, in 0.7% Triton X-100 / 0.1 M HCl (Lab Chem, Cat# LC153004) for 10 min at RT, in 2x SSCT (0.1% Tween-20 in 2x SSC (Promega, Cat# V4261)) for 5 min at RT, in 50% (v/v) formamide (Fisher BioReagents, Cat# BP227-500) in 2x SSCT for 5 min at RT, in 50% formamide in 2x SSCT for 2.5 min at 85°C, and in 50% formamide in 2x SSCT for 20 min at 60°C. Slides were then cooled at RT for 10 min and a 22 mm round coverslip (Fisherbrand, Cat# 12545101) with a primary Oligopaint mix was mounted to the slides and sealed with rubber cement (Elmer’s, Cat# E904). The primary Oligopaint mix was composed of 100 pmol of each Oligopaint, 1.5 µl of 25 µM dNTPs (New England BioLabs, Cat# N0446S), 1 µl molecular grade H_2_O, 12.5 µl formamide, 4 µl PVSA (Aldrich, Cat# 9002-97-5), 1 µl RNase A (VWR Life Science, Cat# E866-5ML), 6.25 µl DNA hybridization buffer (4 g Dextra sulfate sodium salt (Sigma Life Sciences, Cat# 9011-18-1), 40 µl Tween, 4 ml 20x SSC, PVSA up to 10 ml), per reaction. Once the rubber cement had completely dried, the microscope slide was heated to 85°C on a metal block for 2.5 min, and immediately transferred to a 37°C humidified chamber for an overnight incubation. The following day, the cover slips were removed and slides were washed (in coplin jars) in 2x SSCT for 15 min at 60°C, in 2x SSCT for 15 min at RT, and in 0.2x SSC for 10 min at RT. A 22 mm round coverslip with a secondary Oligopaint mix was mounted to the slides and sealed with rubber cement. The secondary Oligopaint mix was composed of 10 pmol of each secondary oligo (IDT, custom synthesized), 6.25 µl DNA hybridization buffer, 12.5 µl formamide, and H_2_O up to 25 µl, per reaction. Slides were then transferred to a 37°C humidified chamber for 2 hr. Following this, the cover slips were removed and slides were washed in 2x SSCT for 15 min at 60°C, in 2x SSCT for 15 min at RT, and in 0.2x SSC for 10 min at RT. A square 22 mm × 22 mm #1.5 coverslip (VWR, Cat# 16004-302) with a drop of Prolong Diamond Antifade Mountain with DAPI (Invitrogen, Cat# P36966) was mounted to the slides. Slides were allowed to set for either 24 hr at RT or 72 hr at 4°C, and then sealed with clear polish.

### Oligopaint FISH of bone marrow cells

To prepare metaphase chromosome spreads, bone marrow from femurs and tibias was flushed out into 2.85 ml of M2 media at 37°C using a 1 cc tuberculin syringe and 23g needle. Bone marrow was then broken up and 150 µl of 0.5% colchicine (Sigma-Aldrich, Cat# C9754-500MG) was added. The suspension was incubated for 10 min at 37°C, and the media was removed by centrifugating at 400 xg for 5 min and removing the supernatant. The pellet was resuspended in 1.5 ml 0.56% KCl solution and incubated for 20 min at 37°C. The suspension was then centrifuged as above, and the supernatant was removed. Cells were fixed with three subsequent rounds of standard washing with methanol/acetic acid (Macron Fine Chemicals, Cat# MK-3016-16 and MK-V193-45) (3/1) fixative solution. The final pellet was diluted in methanol/acetic acid solution and dropped on a clean microscopy slide covered with a steam layer and air dried for two days. To perform Oligopaint FISH, the slides were washed in 1x PBS for 5 min at RT and incubated in 0.005% Pepsin (Sigma-Aldrich, Cat# P7012-250MG) in 0.01 N HCl solution for 10 min at 37 °C. The slides were then washed in 1x PBS at RT and fixed in 1% paraformaldehyde in 1x PBS for 10 min at RT and washed in 1x PBS for 5 min at RT. The slides were dehydrated in a series of graded alcohols (70%, 90%, and 100%), air dried, and the primary Oligopaint mix was mounted to the slides and sealed with rubber cement (Elmer’s, Cat# E904). Subsequent steps were performed as for Oligopaint FISH of oocytes and eggs.

### Microscopy and Image analysis

Fixed oocytes, eggs, and bone marrow cells were imaged with a microscope (Eclipse Ti; Nikon) equipped with 100x / 1.40 NA oil-immersion objective lens, CSU-W1 spinning disk confocal scanner (Yokogawa), ORCA Fusion Digital CMOS camera (Hamamatsu Photonics), and 405, 488, 561 and 640 nm laser lines controlled by the NIS-Elements imaging software (Nikon). Confocal images were acquired as Z-stacks at 0.3 µm intervals. For live imaging, oocytes were placed into 3 µl drops of M2 covered with paraffin oil in a glass-bottom tissue culture dish (fluoroDish, Cat# FD35-100) in a stage top incubator (Tokai Hit) to maintain 37°C. Time-lapse images were collected with a microscope (Eclipse Ti2-E; Nikon) equipped with the 20x / 0.75 NA objective and 60x / 1.40 NA oil-immersion objective, CSU-W1 spinning disk confocal scanner (Yokogawa), ORCA Fusion Digital CMOS camera (Hamamatsu Photonics), and 405, 488, 561 and 640 nm laser lines controlled by the NIS-Elements imaging software (Nikon). For the biased orientation assay in Figure 1D, images of one of the experiments were taken with a microscope (Olympus IX71) equipped with 100x / 1.40 NA oil-immersion objective lens, Visitech VT iSIM scan head, ORCA Quest qCMOS camera (Hamamatsu Photonics), and 405, 442, 488, 514, 561, 640 nm laser lines controlled by MetaMorph acquisition software. Confocal images were collected as Z-stacks at 1 µm intervals to visualize all the chromosomes. Images are displayed as maximum intensity Z-projections. Due to the weak dCas9-GFP signals during live-imaging experiments (Figure 3), we used the Denoise.ai software (Nikon) to reduce noise and follow the *R2d2* locus better in anaphase I. Fiji/ImageJ (NIH) was used to analyze all the images. In general, optical slices containing chromosomes were added to produce a sum intensity Z-projection for pixel intensity quantifications.

### Statistics and reproducibility

Data points were pooled from two to six independent experiments. Data analysis was performed using Microsoft Excel and GraphPad Prism 10. Scattered plots, bar graphs, and line graphs were created with GraphPad Prism 10. Unpaired two-tailed t-test (Figure 2D), chi-square test for goodness of fit for deviations from the expected 50:50 ratio (Figures 2B, 2C, S4, and S5), and chi-square test of independence (Figures 3B and 4B) were used for statistical analyses, and the actual *P* values are shown in each figure or figure legend.

## Supplemental figures

**Figure S1. Oligopaint design to visualize the *R2d2* locus, Related to Figure 1**. (**A**) Oligos were designed to tile the 7 Mb region surrounding the *R2d2* locus (yellow) or the *R2d2* locus itself (green) (See Methods for details). Note that the oligos for *R2d2* locus also recognize the *R2d1* locus because their sequences are highly conserved.^30^ (**B, C**) Bone marrow cells (B) and metaphase I oocytes (C) collected from *R2d2* heterozygous mice were fixed and stained with Oligopaint probes for the 7 Mb region and the *R2d2* locus on chromosome 2.

**Figure S2. dCas9 technique to visualize the *R2d2* locus, Related to Figure 1**. (**A**) Schematic showing how to visualize the *R2d2* locus in mouse oocytes using dCas9. dCas9-EGFP/gRNA complex that recognizes *R2d2* was assembled in a test tube and introduced into prophase I mouse oocytes by microinjection. (**B**) *R2d2* heterozygous (+/–) and control (–/–) oocytes microinjected with dCas9-EGFP/gRNA complex were fixed at metaphase I and stained for EGFP. dCas9-EGFP foci were specifically observed in *R2d2* heterozygous oocytes, confirming that the dCas9 complex recognizes *R2d2*.

**Figure S3. *R2d2* does not induce anaphase lagging by preventing the separation of chromosomes, Related to Figure 3**. *R2d2* heterozygous (+/–) oocytes expressing H2B-mCherry and TALE-mClover2 that specifically recognizes major satellites (centromere) were imaged live at anaphase I.^59^ 96.8% of lagging chromosomes were univalent (n = 31 lagging chromosomes). White line, oocyte cortex. Enlarged insets show a bivalent with two centromere signals and a univalent with one centromere signal.

**Figure S4. Frequent recombination events between the *R2d2* locus and the centromere, Related to Figure 4**. *R2d2* heterozygous (+/–) oocytes were matured *in vitro* and fixed at metaphase II to perform Oligopaint FISH to visualize Chromosome 2 and the *R2d2* locus. The fraction of oocytes with either non-recombinant or recombinant chromosome for the *R2d2* locus was quantified (n = 111 cells); chi-square test for goodness of fit was used to examine the deviation from the expected 50:50 ratio; \**P* < 0.00001; white lines, cell cortex.

**Figure S5. High recombinant rate also in the genetic background without *R2d2* meiotic drive, Related to Figure 4**. *R2d2* heterozygous (+/–) oocytes in the *M. m. domesticus* x *M. m. musculus* hybrid genetic background were matured *in vitro* and fixed at metaphase II to perform Oligopaint FISH to visualize Chromosome 2 and the *R2d2* locus. The fraction of eggs with either non-recombinant or recombinant chromosome for the *R2d2* locus was quantified (n = 107 cells); chi-square test for goodness of fit was used to examine the deviation from the expected 50:50 ratio; \**P* = 0.00016; white lines, cell cortex.

**Table S1. List of primer sets used to produce gRNA targeting the *R2d2* locus.**

**Table S2. List of oligos used for oligopaint FISH in this study**.

