## Supplementary figures and images for "An egg sabotaging mechanism drives non-Mendelian transmission in mice"

### Fig. S1-S5

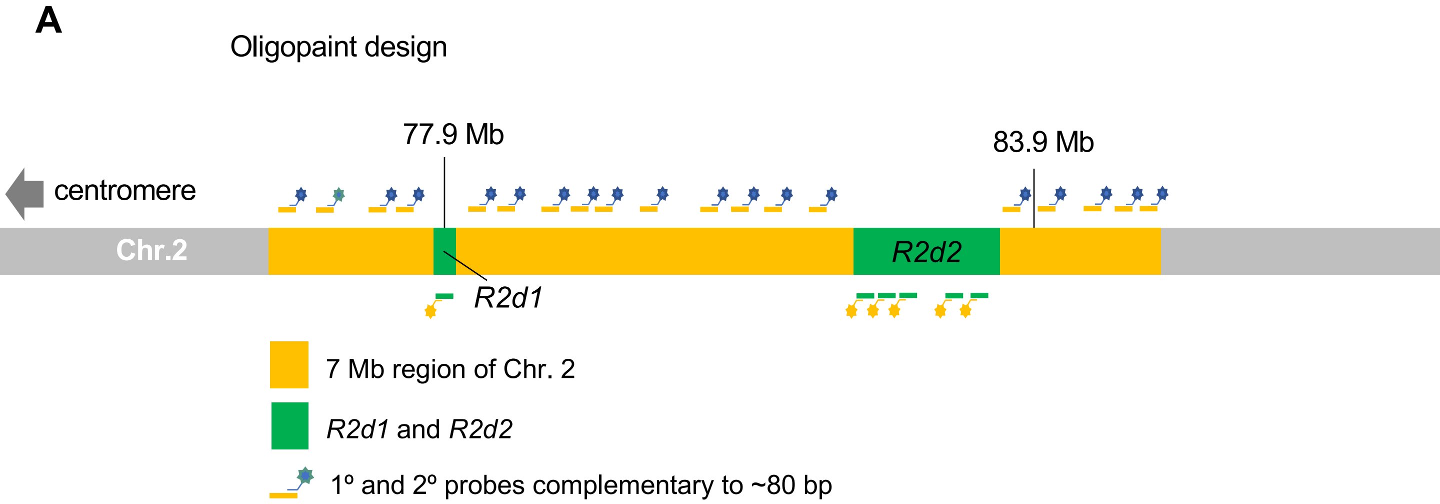

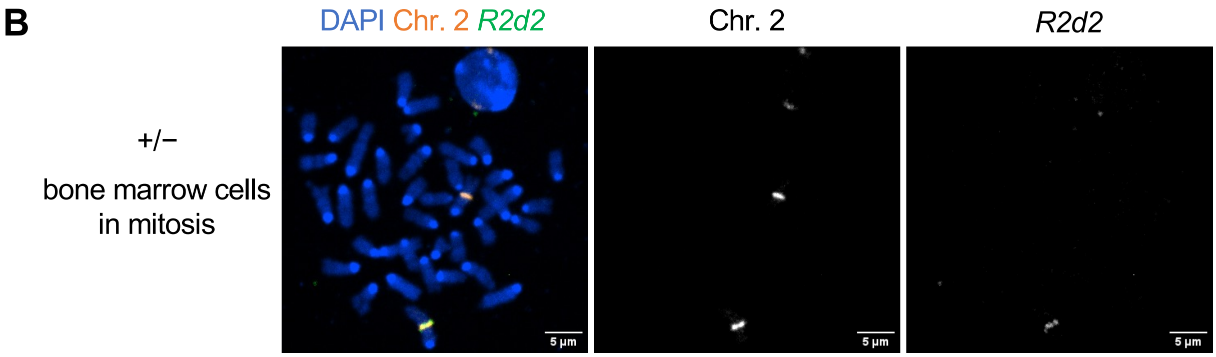

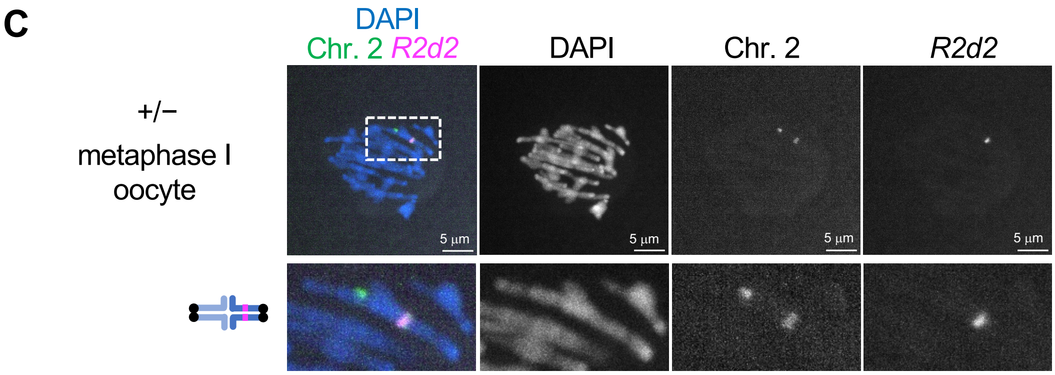

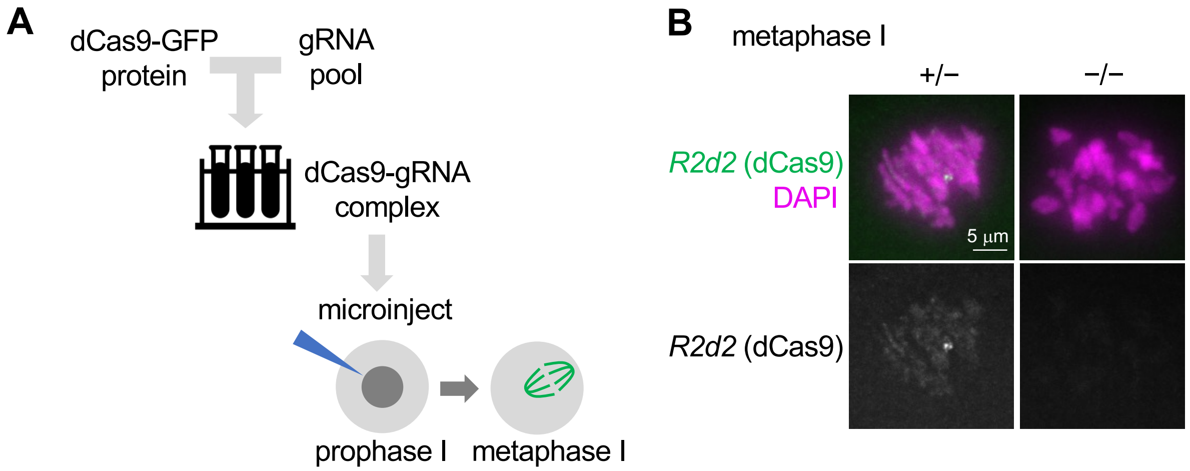

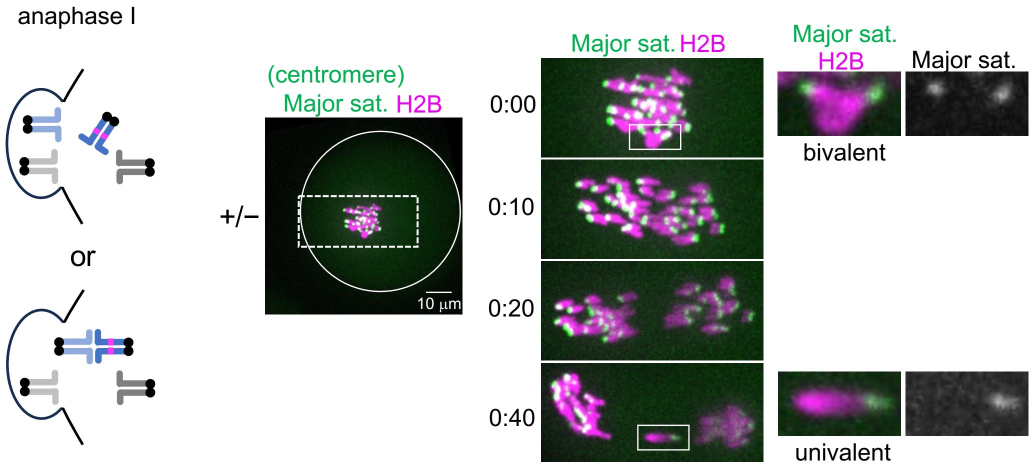

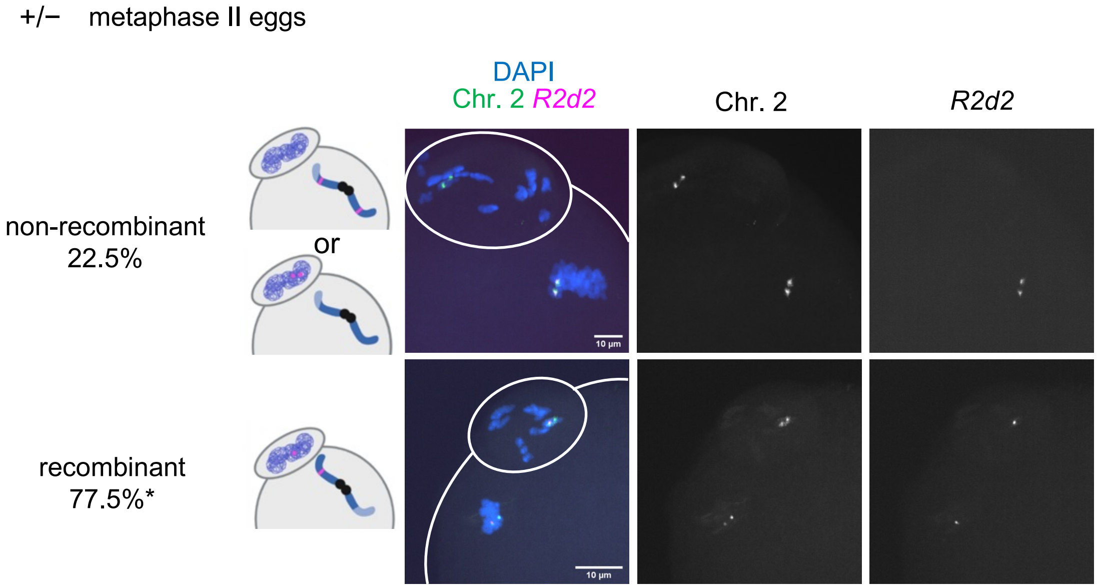

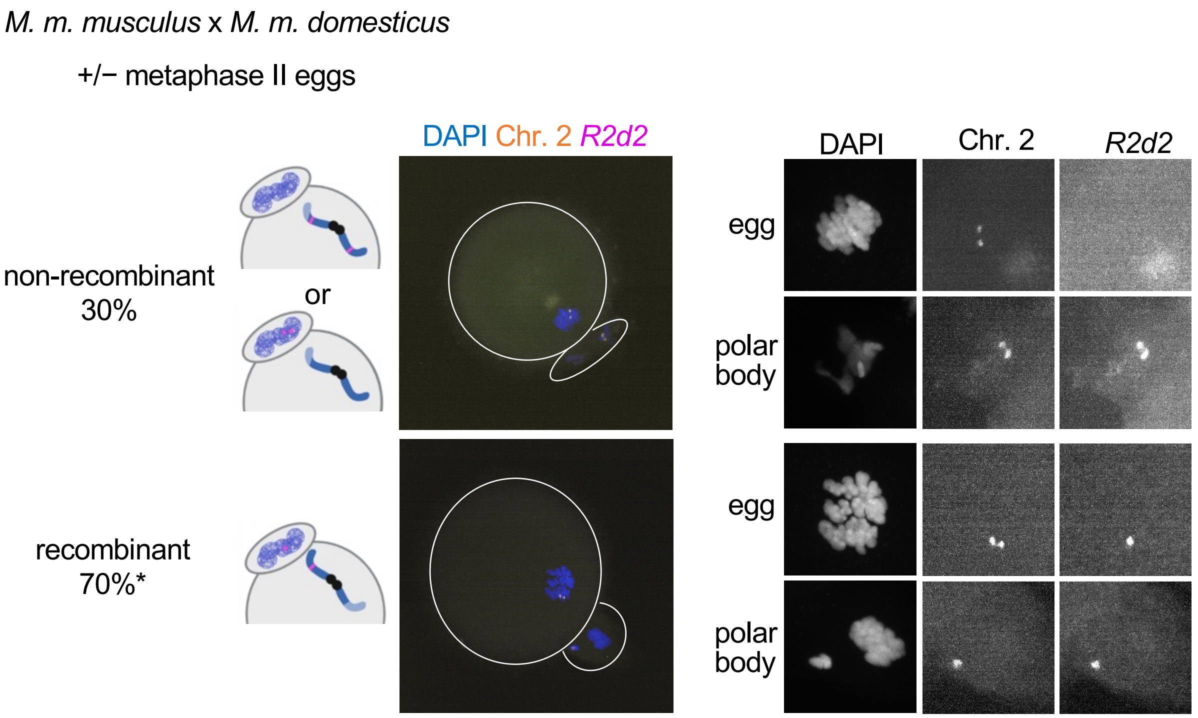
